# Sulfur-species in Zinc-specific Condylar Zones of a Rat Temporomandibular Joint

**DOI:** 10.1101/2024.11.11.623079

**Authors:** Brandon H. Lee, Zhiyuan Yang, Tiffany Ho, Yongmei Wang, Nobumichi Tamura, Samuel Webb, Sharon Bone, Sunita P. Ho

**Author notes:** To whom correspondence should be addressed: Sunita P. Ho, Ph.D., 513 Parnassus Avenue, HSW 813, University of California San Francisco, San Francisco, CA 94143.

## Abstract

In this study, we performed synchrotron-based micro-X-ray fluorescence (μ-XRF) imaging of elements Zn and S, and X-ray absorption near edge spectroscopy (XANES) coupled with μ-XRF for identification of Zn and S species in the condylar zones of a rat temporomandibular joint (TMJ). Histologic localization of Zn and hypoxia-inducible factor-1α (HIF-1α) were mapped using an optical microscope. These data were visually correlated with μ-XRF and XANES data to provide insights into plausible biological S-species in Z-enriched condylar zones of a rat TMJ. Furthermore, μ-XRF coupled with micro-X-ray diffraction (μ-XRD) was used to underline Z-incorporated biological apatite in the subchondral bone and bone of the rat TMJ.

Results illustrated the potential dependence between biometal Zn and nonmetal S and their collective governance of cell and tissue functions in a zone-specific manner. Elemental Zn with organic and inorganic S-species at the cartilage-bone interface and transformation of plausible Zn-enriched mineralization kinetics of biological apatite from subchondral bone to condylar bone were ascertained using μ-XRF-XANES and μ-XRD. The coupled μ-XRF-XANES complementing with μ-XRD and immunohistology provided an informative view of S and Zn and their association with zone-specific biological pathways *in situ*. Understanding the spatial distributions of the main S-species with redox-inert Zn in regions of cartilage, bone, and the interface is essential for further unlocking questions surrounding formation and resorption-related biomineralization pathways as related to osteoarthritis or genetically inherited diseases. Using these complementary techniques with microspectroscopic spatial information provided insights into the associations between biometal Zn and nonmetal S and a window into detecting the plausible early-stage diagnostic biomarkers for humans with TMJ osteoarthritis.

## 1. INTRODUCTION

The compositions of the condylar zones change with mechanical insults on the TMJ (e.g., trauma, iatrogenic dental procedures, and involuntary jaw clenching) (1-6). Chronic mechanical insults are transduced to zone-specific biological pathways that manifest as loss of clinically observed condylar shape and TMJ function. Articular condyles of diarthrodial synovial joints are multizonal biomaterials (7-10) wherein the functional maintenance of each zone is uniquely predisposed to variations in elements, their species, and associations between species (7, 11-17). Spatial associations between elemental zinc (Zn) and sulfur (S) within zone-specific condylar regions can allude to the plausible regulation of the commonly investigated Zn-specific catabolic and anabolic biological pathways (18-20). The individual roles of Zn and S in tissues and their collective partnership in biological pathways continue to be discussed (21-23). Chronic shifts in tissue oxygenation can alter microanatomical levels of the biometal element Zn and the nonmetal element S, postulated to result in ectopic biomineralization within multiple human organs (22-28).

In mineralogy, the mutual affinity of Zn and S is well-recognized (23). However, no studies have addressed the colocalization of Zn and S and their crosstalk to underpin the formation of ectopic biominerals in the human body. This study aims to use microspectroscopic techniques that leverage synchrotron radiation to illustrate the spatial colocalization of Zn with S-species. We will demonstrate predominant organic and inorganic S-species within Z-enriched regions of cartilage, bone, and the hypertrophic and subchondral interfacial regions in a TMJ condyle, with the long-term goal of investigating their collective role in pathobiomineralogy. This first step will help us investigate multifactorial mechanisms that guide downstream ectopic biomineralization events in human TMJ condyles with osteoarthritis.

Microspectroscopy will be performed on rat condylar tissues. Scanning electron microscopy (SEM) will help generate zone-specific cellular and extracellular matrix structure maps. Histologic localization of Zn and hypoxia-inducible factor-1α (HIF-1α, a surrogate for tissue oxygenation) will be mapped using an optical microscope. Micro X-ray fluorescence (μ-XRF) imaging Zn and S elemental colocalization, and X-ray absorption near edge spectroscopy (XANES) coupled with XRF for identification of zone-specific Zn and S species will provide insights into plausible biological pathways in cartilage, cartilage, and bone interface, and bone. The S-species identified using XANES can provide insights into site-specific reduced and oxidized species (14, 29) in condylar zones of the TMJ. Elemental Zn associations with inorganic species at the cartilage-bone interface and condylar bone will be ascertained by micro-X-ray diffraction (μ-XRD) analysis. Using these complementary techniques with microspectroscopic spatial information will allow us to build associations between biometal Zn and nonmetal S and gather insights into early-stage diagnostic biomarkers specifically for humans with temporomandibular joint osteoarthritis (TMJOA).

## 2. MATERIALS AND METHODS

### 2.1. Specimens and tissue sections

Rats often are the small-scale preclinical animal models of choice to investigate the mechanistic pathways of TMJ osteoarthritis (TMJOA). Following the euthanasia of 4-week-old Sprague Dawley rats, the TMJ condyles were surgically extracted (IACUC protocol AN183106-03, UCSF; ARRIVE-Animal Research: Reporting of In Vivo Experiments). Fresh condyles (N=3) were cryopreserved for further examination using XANES. Condyles (N=3) were chemically fixed in 10% neutral-buffered formalin and paraffinized for histology.

### 2.2. Low-energy Microprobe X-ray Fluorescence (μXRF) Imaging

The cryopreserved specimens cut to 8 µm thick sections were mounted on quartz slides for microspectroscopy. Elemental maps specific to Zn, Ca, S, and phosphorus (P) were collected using µ-XRF at the Stanford Synchrotron Radiation Lightsource (SSRL) at beamlines 2-3 and 14-3b. Spatial maps of Zn and Ca were generated at beamline (BL) 2-3 using an incident energy beam of 10keV and a spot size of ∼ 5×5µm. The incident beam energy was selected using a Si(111) monochromator. The fluorescence signal was detected using a 1-element Vortex detector. High-resolution elemental maps were acquired at regions of interest with a spot size of ∼ 1×1µm. Spatial maps of elements P and S were generated at BL14-3b using an incident energy beam of 2488eV and a spot size of ∼ 5×5µm (incident beam energy selected using a Si(111) monochromator). Multi-energy maps of S were acquired at 2472.68, 2473.27, 2476.2, 2482.4, 2482.7eV to map glutathione disulfide, glutathione/methionine, sulfoxides, organic sulfates, and inorganic sulfates, respectively using a 7-element Vortex detector. The monochromator was calibrated by setting the maximum of the first peak of a Na_2_S_2_O_3_ spectrum to 2472.02 eV. Data were analyzed using Sam’s Microprobe Analysis Kit (SMAK) (30, 31).

### 2.3. X-ray Absorbance Near-Edge Spectroscopy

Zn K-edge XANES spectroscopy was performed at points within regions of interest in tissues. Points on condylar zones were selected using the Zn XRF map as a guide. Zn spectra were acquired from higher/lower Zn regions from energy levels 9640 to 9760eV with a spot size of ∼ 1×1µm at BL 2-3. Spectra were calibrated using a Zn foil and selected the maximum of the first derivative to be 9659.0 eV.

Multi-energy S XRF maps were used to create maps of variance by using algorithms in SMAK (30, 31). Points of interest for S-spot XANES were selected by anatomical locations, higher/lower regions of S, and spots with higher/lower variance. S spectrums were acquired from 2465eV to 2490eV with a spot size of ∼ 5×5µm at BL2-3. Data were background corrected and fit to in-house Zn and S standards using Sam’s Interface for XAS Package (SIXPack) (32).

### 2.4. Micro-X-ray Diffraction Analysis

Synchrotron 2D micro-X-ray diffraction (µXRD) patterns were collected using BL12.3.2 of the Advanced Light Source at Lawrence Berkeley National Laboratory. Lower-resolution Zn and Ca XRF maps were used to locate regions of interest. X-ray diffraction patterns were measured using a monochromator with an energy of 10keV, wavelength λ=1.2398Å, and reflective geometry. The area detector (DECTRIS Pilatus 1M) was placed at an angle of 40° at 158mm from the sample surface with an X-ray beam spot of 10×3μm. Data were analyzed using in-house software XRDSol and fit to hydroxylapatite standards acquired from the MINCRYST database (33).

### 2.5. Field emission scanning electron microscopy

Following beamline spectroscopy, the quartz-mounted section was mapped using field emission scanning electron microscopy (FESEM) (SIGMA VP500, Carl Zeiss Microscopy, Pleasanton, CA). Surface morphologies were imaged at an incident electron energy of 1keV.

### 2.6. Histology

The NBF-fixed specimens were embedded in paraffin, and 5µm thick sections were mounted on glass slides. Sections were deparaffinized and rehydrated, before staining with 0.1% Safranin-O for 5min. Counter-staining was achieved with 0.001% Fast Green for 5min (34, 35). Images were acquired using a brightfield microscope (AxioObserver2, Carl Zeiss Microscopy, Pleasanton, CA).

### 2.7. Immunohistology

Enzymatic epitope retrieval for immunohistochemistry was achieved by treating tissue sections with HistoReveal (ab103720, Abcam) at room temperature for 10min. Blocking was performed with a commercial hydrogen peroxidase block (ab236469, Abcam, Waltham, MA) for 10min and 1% BSA/PBS for 5 min. The primary antibody for HIF-1α (100:1, Abcam, Waltham, MA) was diluted in 1% BSA/TBS and incubated under static conditions overnight at 4°C. Further incubation with HRP-conjugate occurred at room temperature for 15 min (ab236469, Abcam, Waltham, MA). The section was treated with a secondary antibody (Alexa Fluor 555, Abcam, Waltham, MA) at room temperature for 1hr, co-localized with Zn using 5µm FluoZin™-3 - tetrapotassium salt (F24194, ThermoFisher, San Francisco, CA) stained at room temperature for 1hr, and counter-stained with DAPI (D1306, ThermoFisher, San Francisco, CA). Images were acquired using fluorescence microscopy (AxioObserver2, Carl Zeiss Microscopy, Pleasanton, CA).

## 3. RESULTS

Microscopy and spectroscopy datasets on physical, biochemical, and elemental characteristics across condylar zones from serial sections of the same condyle were visually correlated. The red asterisk illustrates a common feature across all tissue sections used in this study (**Figs. 1-3**).

**Figure 1.**
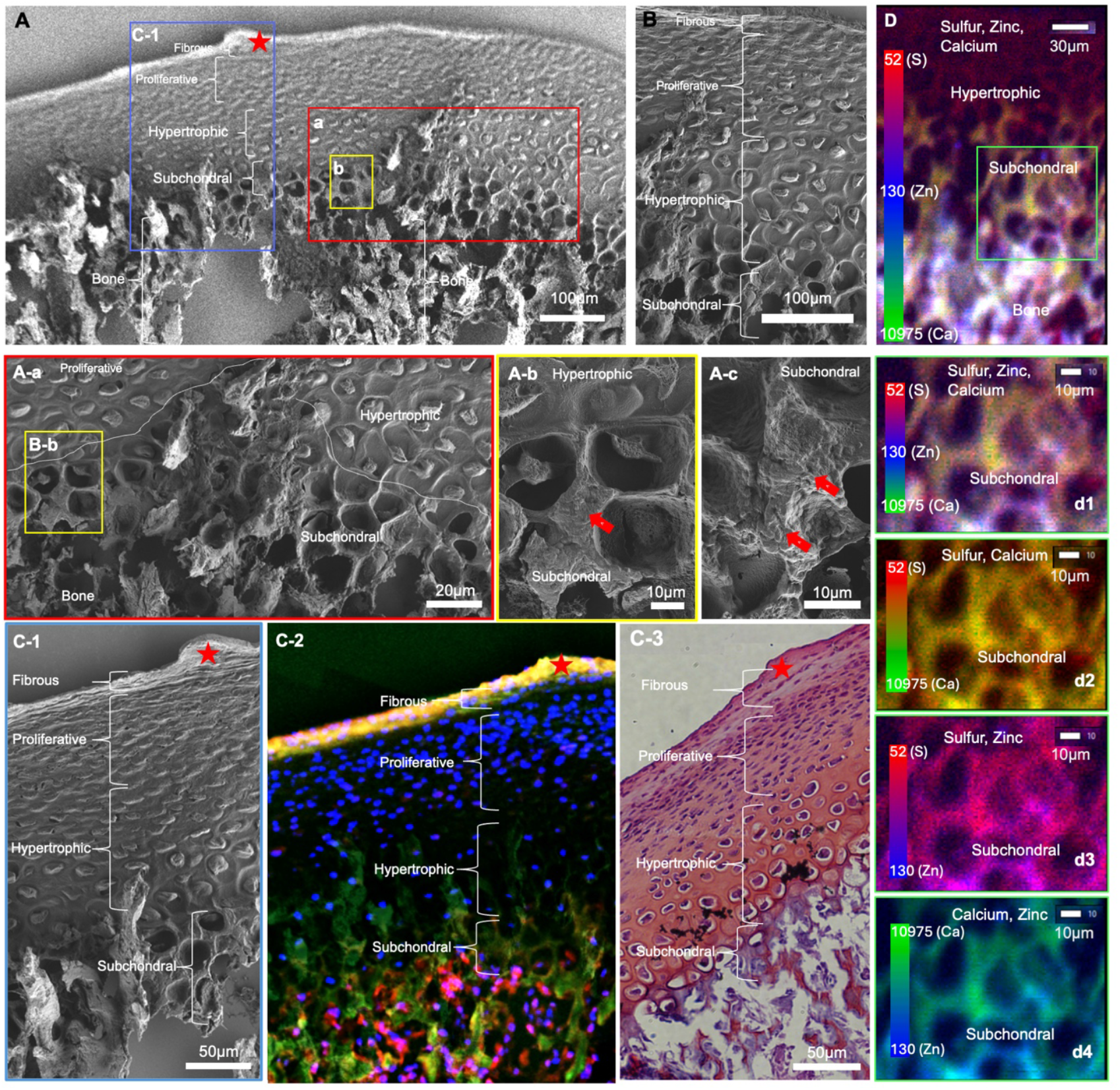
Zonal classification of cartilage, cartilage-bone interface, and bone: **A**. Zone-specific structure of a tissue section from a rat condyle using a scanning electron micrograph. Regions a-c illustrate zone-specific cell morphology at the cartilage-bone interface **(A-a)** (redlined rectangle), hypertrophic (yellow-lined rectangle), and subchondral regions **(A-b)**. Subchondral regions at a higher magnification illustrate spherical nodules in the perilacunar walls (yellow-lined rectangle, arrows – **A-b, A-c**). **C**. Zone-specific correlation between structures (**C-1**), zinc (Zn) and HIF1α **(C-2)**, and Safranin O-stained sulfur (S)-rich proteoglycans **(C-3). D**. X-ray fluorescence of zone-specific S, Zn, and calcium (Ca) elements with all three elements colocalized in the bone and to a lesser extent in the subchondral bone (**Supplemental Figure 1)**.

### 3.1. Structural, elemental, and histological compartmentalization of condylar zones; fibrous, proliferative, hypertrophic, subchondral, and bone

SEM of condylar zones (**Figs. 1A-a, A-b, A-c)** at higher magnification illustrated morphologically distinct cells at the hypertrophic (**Fig. 1A-a**), and subchondral regions **(Figs. 1A-a, 1A-b)**. Subchondral regions at a higher magnification illustrated spherical nodules in the perilacunar walls (red block arrows in **Figs. 1A-b**, and **1A-c**). Zone-specific correlation between structure and localized Zn with HIF1α **(Fig. 1C-2)**, and visually correlated with Safranin O **(Fig. 1C-3)** indicated an association between Zn-HIF1α-PGs in the fibrous and subchondral regions, and Zn and PGs in the hypertrophic zone. No visual association between Zn and PGs was apparent in the proliferative zone (**Figs. 1C-2, 1C-3**). X-ray fluorescence of zone-specific S, Zn, P, and Ca elements were colocalized in the bone and to a lesser extent in the subchondral bone (**Fig. 1D, Supplemental Fig. 1**). Visual correlation between spatially immunolocalized biomolecules and multi-energy XRF maps confirmed the colocalization of plausible Z- and S-species in the hypertrophic and subchondral zones of cartilage, and bone, but not in fibrous zone of cartilage (**Figs. 1-3**).

### 3.2. Sulfur species in condylar zones

The coupled μ-XRF and XANES method detected specific S species in a rat condyle at high spatial resolution. Individual multi-energy XRF maps illustrated zone-specific localization of S species glutathione disulfide (GSSG), glutathione (GSH), sulfoxides, organic sulfates, and inorganic sulfates within different condylar zones and bone (**Fig. 2A**). Individual spot S-XANES from cartilage, bone, and the subchondral-bone interface (**Supplemental Fig. 2**) and average S XANES for cartilage (ii), interface (iii), bone (iv), and S standards (v) are shown. Dotted lines correspond to energies at which multi-energy maps (**Fig. 2A**) of tissue sections were acquired and correspond to peaks in the spectra of the different standards. Results illustrated the ability to map S species and establish their zone-specific colocalization, such as GSH (or methionine), and their oxidized form GSSG and methionine sulfoxide (2475.9eV) (29), as well as organic sulfate as represented by chondroitin sulfate proteoglycan (PG) (2482.4eV) (31) or more generally sulfate ester (as chondroitin sulfate PG at 2482.7eV) (31). Sulfoxide distributions were visually correlated with GSH/methionine and GSSG and could be a part of methionine sulfoxide (**Fig. 2A-C**).

**Figure 2.**
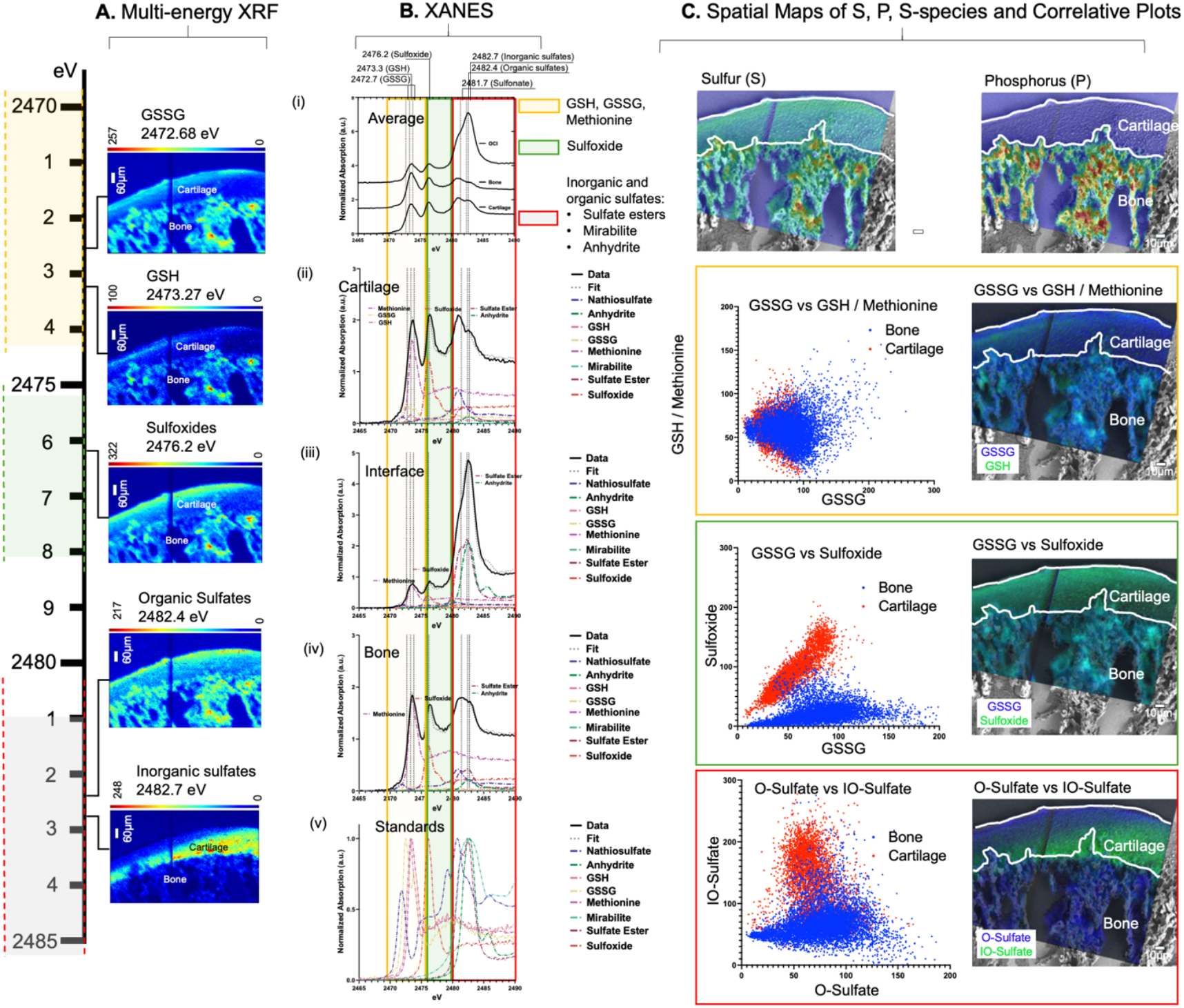
Sulfur species in cartilage and bone: **A**. Individual multi-energy XRF maps illustrate zone-specific localization of sulfur (S) species GSSG, GSH, sulfoxides, sulfonates, and organic and inorganic sulfates across a rat condyle (scale bars represent 60µm and color bars represent counts). **B.(i)**. Averaged spectra of all S XANES points from cartilage, bone, and the subchondral-bone interface (**Supplemental Figure 2**) are shown. Individual but average S XANES **(i)** for cartilage **(ii)**, interface **(iii)**, bone **(iv)**, and S standards **(v)** are shown. Dotted lines correspond to peaks in the spectra from respective standards at which multi-energy maps (column A) of the tissue section were acquired. **C**. Row 1 – XRF counts of S and phosphorus (P) in cartilage and bone are overlayed on the matrix structure. As expected, S can be observed in cartilage and to a lesser extent at the interface but higher in bone. Row 2 – Visual overlays of the bicolor plot with matrix structure show the colocalization of respective S species in condylar regions (right) (scale bar is 10μm). The corresponding correlation plots of the S species are shown to the left of the overlays.

The sulfate esters identified by XANES spectroscopy could be representative of the sulfated PGs in cartilage and bone (31). It is likely that the co-localization of sulfoxides with disulfide glutathione (GSSG) using multi-energy μXRF mapping and the observation of methionine using via XANES (**Supplemental Fig. 2**) indicates methionine sulfoxide are an additional potential species within the cartilage, bone, and its cartilage-bone interface. However, in cluster maps, the trend between GSH and GSSG appeared to be similar for cartilage and bone (**Fig. 2C**). This is likely because methionine is similar across cartilage and bone and makes up the majority of the signal with only minor contributions from GSH and GSSG (Fig. 2B). This interpretation of XANES signal can be further ascertained by performing “XANES fitting” of the multi-energy maps (31). As mentioned earlier, GSSG is a minor species compared to methionine (Fig. 2B). The methionine-sulfoxide could be the dominating compared to the GSSG-sulfoxide relationship (**Fig. 2C**). Based on these results, methionine and methionine sulfoxide (2473.1 and 2475.9eV) could likely be used to identify the extent of oxidative stress in cartilage, while the ratio between GSH and GSSG (2473 and 2472.2eV) can be used for bone.

### 3.3. Zn-species and Zn incorporation in apatite lattice of condylar zones

The Zn μ-XRF map overlaid on the SEM micrograph enabled visualization of Zn in the subchondral regions and bone. These observations (**Fig. 3A**) agreed with the enzymatically localized Zn in the same regions (**Fig. 1C-2**). At a 5 μm spatial resolution, Zn in the fibrous layers was not detected using μ-XRF. At a 1μm spatial resolution, the signal-to-noise ratio of Zn in the fibrous layer of cartilage was low. However, detectable Zn levels in the hypertrophic and subchondral condylar regions and bone from BL 2-3 (**Fig. 3B**) were confirmed using XRF with a 5μm spot size. As mentioned, the mass of the tissue to interaction volume with the beam could be different between cartilage and bone. Future experiments should likely consider region-specific section thickness and XANES acquisition parameters.

**Figure 3.**
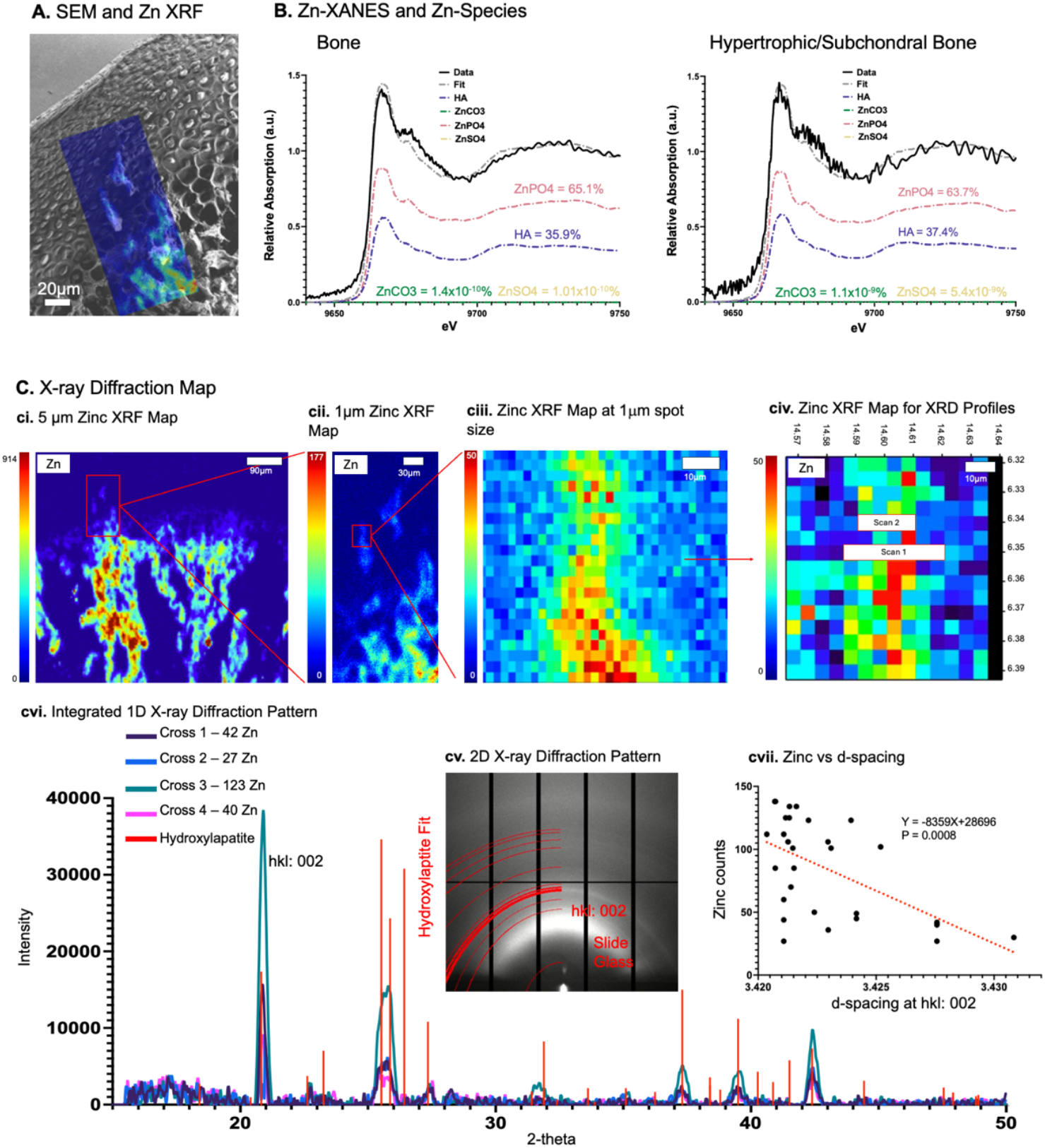
Zinc species and Zinc incorporation in apatite lattice in the bone-cartilage interface: **A**. XRF map of zinc (Zn) overlaid on tissue structure to visualize Zn localization in subchondral regions and bone is in agreement with enzymatically localized Zn (**Fig. 1C-2**). **B**. Zn XANES and plausible Zn coordination with other nonmetal elements in cartilage and bone include nitrogen, sulfur (S), and phosphorus. Species could include Zn carbonates, sulfates, phosphates, and nitrides. **C**. Left to right – Detectable Zn levels at 5µm spot size from SSRL BL 2-3 illustrate its presence in subchondral and bone regions of the rat condyle (ci). At 1µm spot size, the Zn levels within the red rectangle can be observed in detail (**ciii**). The average XRD patterns from scan lines 1 and 2 (scan 1 and 2, **civ**) are shown (**cv**), and the 2theta vs. intensity is shown in **cvi**. Zn counts vs. d-spacing hkl (002) illustrated smaller d-spacing with increased Zn counts (**cvii**).

Plausible Zn species such as phosphates and carbonates in hypertrophic cartilage and subchondral bone, and bone using XANES are shown in **Fig. 3B** at a 5μm spot size. At 1µm spot size details in Zn levels were identified within the subchondral regions (**Figs. 3cii, 3ciii**). The XRD scan lines 1 and 2 (**Fig. 3civ**) from the Zn positive regions within the red rectangle (**Fig. 3cii**) illustrated an XRD pattern for hydroxylapatite (**Fig. 3cv**) with decreased d-spacing hkl (002) for increasing Zn counts (**Fig. 3cvii**).

## 4. DISCUSSION

The motivation for this study is to investigate compositional changes in the condylar zones as potential early-stage diagnostic biomarkers of TMJOA. The zonal compartmentalization of S-rich condylar cartilage is elaborated as differences in collagen orientation, PG concentration, cellular morphology, water content, and mineralization (7, 11, 34-38). These noted downstream differences are postulated from upstream changes in intra-and extra-cellular biochemical expressions that regulate anabolic and catabolic biological pathways necessary for maintaining cartilage, bone, and the interface.

Visual correlation between spatially immunolocalized biomolecules, multi-energy XRF maps, and XANES confirmed the colocalization of plausible Z- and S-species in condylar zones (**Figs. 1-3**). Zone-specific Zn with hypoxia factors (HIF-1α) in a matrix rich with **1)** fibrous articular cells (**Fig. 1**) and organic sulfates, including reduced (GSH/methionine) and oxidized (GSSG/sulfoxides) forms, sodium thiosulfates and sulfate-esters (sodium thiosulfates as plausible sodium chondroitin sulfated complexes (39-41), sulfate esters as possibly chondroitin sulfated PGs - (31)); **2)** hypertrophic and subchondral cells with inorganic sulfates with sulfate-esters as salts such as mirabilite and anhydrite (42) were spatially correlated (**Fig. 2**).

The observed Zn and HIF-1α in the fibrous regions of cartilage, hypertrophic and subchondral bone are positive regulators of collagen and sulfated PGs and promote chondrocyte differentiation and matrix synthesis necessary for the health and function of articular cells of the fibrous cartilage, and hypertrophic cells in subchondral bone that are in direct proximity with osteoblasts and osteoclasts in bone (17, 19, 43, 44). Standard histologic stained regions by Safranin O, a cationic dye that is sensitive to the polyanionic sulfated PGs (45), are comparable to μ-XRF of S in condylar zones (**Fig. 1**). The PG function generally is regulated by sulfation. Sulfates are added to specific sugar residues as esters to maintain tissue integrity and growth factor signaling necessary for tissue development and its maintenance. From a biomechanics standpoint, the S-rich PGs are chemical and structural “tacks” that bind water, maintain collagen fibrillar spacing, and provide the overall structure and function of the extracellular matrix. Sulfation in the cartilage matrix is directly proportional to bound water and determines the compression resistance of the multizonal cartilage (46). Spectroscopically observed higher sulfate esters possibly from sulfated PGs and methionine from the ECM of cartilage (**Figs. 2B, 4C**) could be the contributors to the observed organic sulfate μ-XRF signal (**Fig. 2A**) in the Zn and HIF-rich articular cartilage zone (**Fig. 1C2**). Spectroscopically observed lower sulfation (sulfate esters) and methionine, and higher levels of sodium thiosulfate, anhydrite, and mirabilite (**Figs. 2B, 4C**) could be the contributors of the observed organic and inorganic sulfates (Figs. 2A) in the Zn and HIF-rich hypertrophic and subchondral zones (**Fig. 1C2**). The cells in the Zn-replete proliferative mid-zone (**Fig. 1C2**) are likely coaxed along the differentiation pathway to regulate chondrocyte hypertrophy (47). Hypertrophic chondrocytes promote changes in surrounding tissue, increase Zn-enriched minerals and bone replacement, and subsequently undergo apoptosis to maintain joint homeostasis (79) (**Fig. 2**). The observed nodular aggregation reinforcing the organic perilacunar walls within this region was akin to matrix vesicle-mediated endochondral ossification observed in our previous studies (48). Observed inorganic sulfates (**Figs. 2A, 2B**) are chelators of metal ions, and in this study possibly of Zn (49). Both organic and inorganic sulfates exist in the presence of Zn and allude to its multi-functional role in the maintenance of the articular layer of cartilage and the subchondral interface with bone. The co-localization of S, Zn, and Ca in the bone and to a lesser extent in the subchondral bone (**Fig. 1D, Supplemental Fig. 1**), hints at the transformation of plausible Zn-enriched mineralization kinetics of apatite from subchondral bone to bone. Given all this information, it is conceivable that Zn can co-exist with organic and inorganic sulfates in the multizonal TMJ condyle.

Based on biomolecular (histology) and elemental (XRF) distributions of Zn and knowledge gained on zone-specific S-species from multi-energy XRF and spot XANES, we can infer that Zn coordinates with sulfates in multiple condylar zones (Figs. 1-3). Based on Zn XRF maps, spot-specific Zn XANES, we can however confirm carbonates and phosphates in both subchondral bone and bone (**Fig. 4**). In this study, more importantly, a nuanced signal highlighting the multiple roles of biometal Zn at the interface and in association with S salts in the presence of apatite was identified for the first time. Spectroscopically, methionine and sulfate ester dominated in the Zn-enriched fibrous region of cartilage and mineralized bone compared to the anhydrite and mirabilite Zn-enriched region of the subchondral interface (**Fig. 4**). In bone, sulfate ester, and methionine as organic sulfates and traces of sodium thiosulfate as inorganic sulfate could be the contributors. Although Zn-carbonate and -phosphate likely have representative X-ray diffraction patterns, in this study, the diffraction pattern was dominated by hkl index of the 002-lattice plane of calcium-based apatite. Decreased d-lattice spacing, specifically at higher Zn counts, indicated its incorporation into a hydroxylapatite lattice structure (50-52). Different anionic/cationic substitutions and vacancies are generally expressed in biological apatite, and conceivably Zn substitution within the lattice structure should be no surprise.

**Figure 4.**
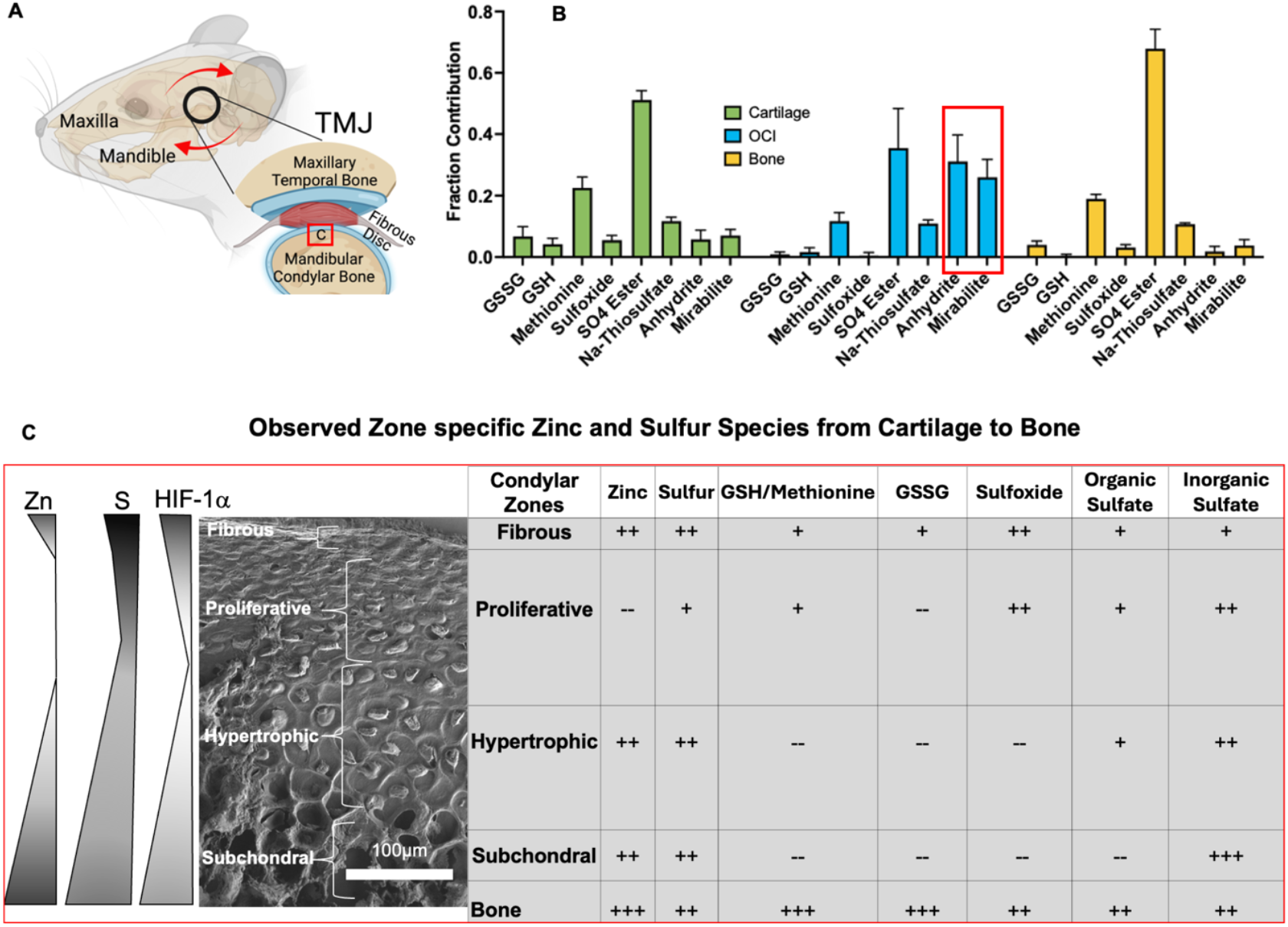
Summary of zone-specific elemental and species localization in cartilage, interface, and bone. **A**. Schematic showing the location of the TMJ and the mandibular condyle. **B, C**. Observed zone-specific semiquantitative measures of Zn- and S-species from cartilage to bone are summarized.

In summary, the S species in Zn-enriched regions of the cartilage and bone and the interface can be characterized using high-resolution microspectroscopy techniques to provide insights into plausible Zn-bound S species as potential biomarkers to detect early-stage pathology in the biomechanically active TMJ condyle. The Zn and S species were mapped to underpin zone-specific oxidative risk in cartilage zones of a rat condyle. Results from this study also hint at the potential dependence between biometal Zn and nonmetal S and their collective governance of cell and tissue functions in a zone-specific manner. We expect that this anatomy-specific Zn coordination chemistry changes with the extent of mechanical insult on the TMJ over time. The coupled μ-XRF and μ-XANES approach allowed for an informative view of S and Zn and their association with biological systems *in situ*. A further extension of the data could be XANES fitting of multi-energy maps (31, 53) collected across the S-K edge to confirm the observed S-species and distinguish these species and their distributions in Zn-enriched condylar regions.

Understanding the spatial distributions of the main S-species with redox-inert Zn in regions of cartilage, bone, and the interface is essential for further unlocking questions surrounding formation and resorption-related biomineralization pathways as related to osteoarthritis or diseases manifested through genetic inheritance. A logical extension as a future study would be to compare and contrast S- and Zn-species levels in condyles under chronic oxidative stress, a postulated insult for temporomandibular joint osteoarthritis.

## Supporting information

All Supplemental Materials

## ACKNOWLEDGEMENTS

This research used beamline 12.3.2 of the Advanced Light Source, a DOE Office of Science User Facility under contract no. DE-AC02-05CH11231. Use of the Stanford Synchrotron Radiation Lightsource, SLAC National Accelerator Laboratory, is supported by the U.S. Department of Energy, Office of Science, Office of Basic Energy Sciences under Contract No. DE-AC02-76SF00515. The SSRL Structural Molecular Biology Program is supported by the DOE Office of Biological and Environmental Research, and by the National Institutes of Health, National Institute of General Medical Sciences (P30GM133894). The contents of this publication are solely the responsibility of the authors and do not necessarily represent the official views of NIGMS or NIH. The image of the rat TMJ was created with BioRender.com.

## AUTHOR CONTRIBUTIONS

**Brandon Lee:** contributed to the experimental setup, synchrotron data acquisition and analyses from BLs 2-3 and 14-3, and data interpretation; **Zhiyuan Yang, Tiffany J. Ho, and Yongmei Wang:** contributed to data acquisition from rat tissues and performed histology; **Nobumuchi Tamura:** Acquired XRF/XRD data and critically reviewed data, and provided data interpretation, and edited the manuscript; **Samuel Webb and Sharon Bone:** assisted with μXRF, critically reviewed zinc and sulfur XANES, provided data analysis and interpretation, and edited the manuscript; **Sunita P. Ho:** contributed to concept and study design, data analysis and interpretation, drafted the manuscript, revised and critically reviewed the manuscript with all authors. All authors gave final approval and agreed to be accountable for all aspects of the work.

## DATA AVAILABILITY STATEMENT

The data that support the findings of this study are available from the corresponding author upon reasonable request.

## FIGURE LEGENDS

**Supplemental Figure 1 (Fig. S1)**. Spatial colocalization (A) and cluster plots (B) of sulfur (S) and zinc (Zn), calcium (Ca) and zinc (Zn), and calcium (Ca) and phosphorus (P). Cluster plots illustrate a narrow window of variation in Zn for a large variation in S in cartilage. The overlap in Ca and Zn, and Ca and P of cartilage with bone is predominantly from the hypertrophic and subchondral regions within cartilage and the interface with bone.

**Supplemental Figure 2 (Fig. S2): A**. Sparse principal component analysis (sPCA) and simplex volume maximization (SiVM) of sulfur (S) ME maps. Principal components sPCA1-4 illustrate sites of variance. **B**. XANES points (1-11) were chosen based on anatomical location (i.e. cartilage, interface, bone), S counts (i.e. high/low), and variance (i.e. high/low) (B). **C**. All XANES curves are organized by region (C). Point 9 was removed because of low signal-to-noise ratio.

**Supplemental Figure 3 (Fig. S3):** Energies at which sulfur (S) species are typically detected by various researchers using X-ray absorption near edge spectroscopy (XANES).

