## Supplementary material for "Sulfur-species in Zinc-specific Condylar Zones of a Rat Temporomandibular Joint": All Supplemental Materials

[1]Preventive and Restorative Dent. Sci., [2]Urology, University of California, San Francisco, San Francisco, CA; [3]Neuroscience Graduate Group, University of California, Davis, Davis, CA; [4]School of Dentistry, University of Washington, Seattle, WA; [5]Oral and Maxillofacial Surgery, Veterans Affairs San Francisco Health Care, San Francisco, CA; [6]Advanced Light Source, Lawrence Berkeley Natl. Lab., Berkeley, CA; [7]Stanford Synchrotron Radiation Lightsource, SLAC National Accelerator Laboratory, Menlo Park, CA

*To whom correspondence should be addressed:

Sunita P. Ho, Ph.D.
513 Parnassus Avenue, HSW 813
University of California San Francisco
San Francisco, CA 94143

**FIGURE LEGENDS**

**Supplemental Figure 1 (Fig. S1).** Spatial colocalization (A) and cluster plots (B) of sulfur (S) and zinc (Zn), calcium (Ca) and zinc (Zn), and calcium (Ca) and phosphorus (P). Cluster plots illustrate a narrow window of variation in Zn for a large variation in S in cartilage. The overlap in Ca and Zn, and Ca and P of cartilage with bone is predominantly from the hypertrophic and subchondral regions within cartilage and the interface with bone.

**Supplemental Figure 2 (Fig. S2): A.** Sparse principal component analysis (sPCA) and simplex volume maximization (SiVM) of sulfur (S) ME maps. Principal components sPCA1-4 illustrate sites of variance. **B.** XANES points (1-11) were chosen based on anatomical location (i.e. cartilage, interface, bone), S counts (i.e. high/low), and variance (i.e. high/low) (B). **C.** All XANES curves are organized by region (C). Point 9 was removed because of low signal-to-noise ratio.

**Supplemental Figure 3 (Fig. S3):** Energies at which sulfur (S) species are typically detected by various researchers using X-ray absorption near edge spectroscopy (XANES).

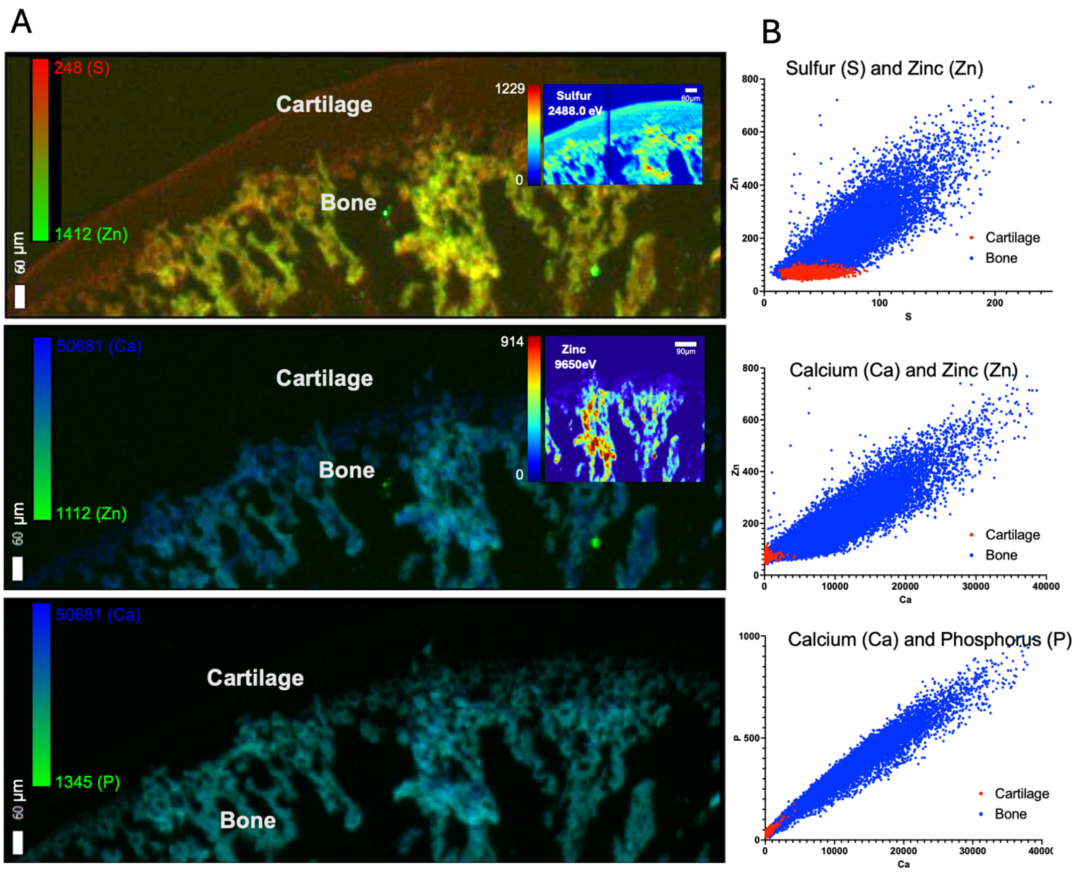


Supplemental Figure 1

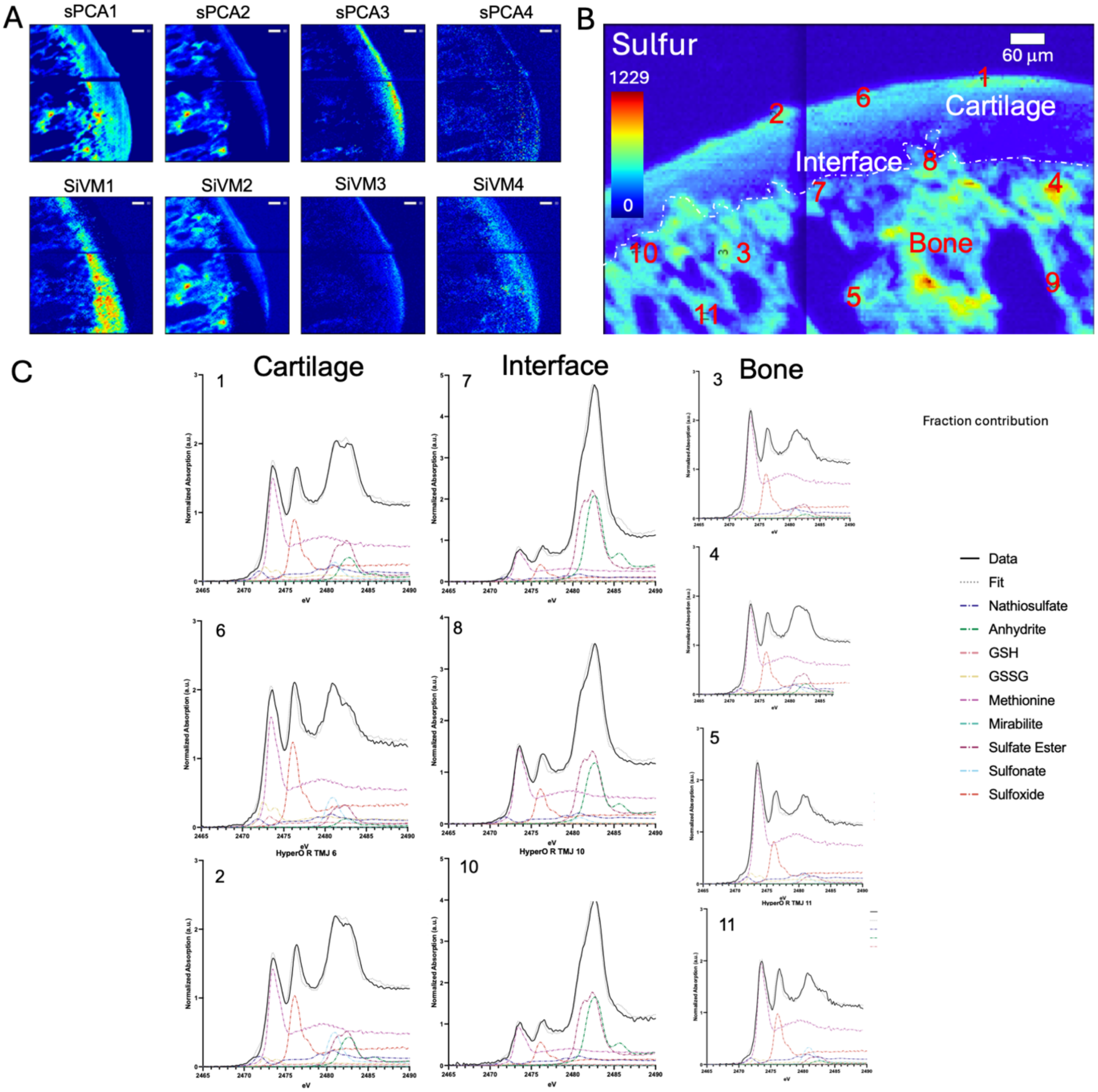


Supplemental Figure 2

| References for S-species using X-ray absorption near edge spectroscopy (XANES) | | | | | | |
|---|---|---|---|---|---|---|
| Reference/S-species | GSSG | GSH | Sulfoxide | Sulfonate | Organic Sulfate | Inorganic Sulfate |
| Our study | 2472.68 | 2473.27 | 2476.2 | 2481.68 | 2482.4 | 2482.7 |
| XANES Standards | 2472.5 | 2473.25 | 2476.1 | 2481 | 2482.35 | 2482.65 |
| Analytical Chemistry. 2018;90(21):12559-66 | 2472.8<br>2473.9 | n/a | 2476.1 | 2481.4 | 2482.4 | 2482.7 |
| ESRF | 2472.6 | 2473.4 | 2476.3 (dmso) | 2481.0 (taurine) | 2482.7 (chondroitin sulfate) | 2482.6 (anhydrite) |

Supplemental Table 1
